# Urban soil microbiomes exhibit taxonomic and functional potential for enhanced contaminant cycling

**DOI:** 10.1101/2024.05.09.592449

**Authors:** Izabel L. Stohel, Young C. Song, Anna B. Turetcaia, Andrew J. Wilson, Dietrich Epp Schmidt, Stephanie A. Yarwood, Andrew Townsend, Emily B. Graham

## Abstract

Urbanization is a leading cause of global biodiversity loss, but its impact on soil microorganisms and biogeochemistry remain uncertain. Although all urban soil microbiomes are influenced by common anthropogenic processes, we lack generalizable patterns in how urbanization impacts the composition and functional potential of the global soil microbiome. These patterns are vital for grasping cities’ roles in biogeochemical processes and managing urban land expansion. We re-analyze soil metagenomic sequences from the Global Urban Soil Environment Ecology Network (GLUSEEN) to identify coordinated changes in microbial taxonomy and functionality and to pinpoint core taxa and metabolisms across five global cities spanning ecoregions and management regimes. Our findings reveal greater spatial variability in microbial taxonomy compared to functional potential, indicating that common anthropogenic processes might shape the genetic potential of diverse taxa. We identify 111 gene annotations that represent core urban soil functions, present in all cities but rare (<5%) in reference soils. Across low-to-high urban land use gradients; nitrogen, trace/heavy metal, glycerol, and fructose/fructan metabolic processes are linked with highly urbanized areas. Core urban microbial taxa – defined by overrepresentation in urban land uses compared to reference soils – included diverse soil bacteria and a more constrained set of methane- and nitrogen-cycling archaea. Overall, our work aids in deriving testable hypotheses and in modeling the feedbacks between growing metropolitan areas and the processes impacting global environmental change.

## Introduction

Despite urban areas covering only 1% of global land, 70% of the world’s population is projected to live in cities by 2050 (United Nations 2018; Ritchie, Samborska and Roser 2024). Human activity in these urban centers is reshaping the way organisms evolve and interact with each other (Pickett *et al*. 2011). Research on urbanization’s effect on ecosystems is becoming more common, but knowledge about urban soil microbiomes is still sparse, even though they play a crucial role in global biogeochemistry (Dubey *et al*. 2019; Naylor *et al*. 2020). Therefore, a comprehensive understanding of soil microbial community structure and functional potential is essential for predicting the ecological roles of microbes in urban ecosystems.

Rapidly increasing urbanization and associated changes in land-use are responsible for biodiversity loss of macroorganisms (i.e., animal and plant communities) and for increased pollution, both of which may subsequently impact urban biogeochemistry and soil microbiome function (Pauchard *et al*. 2006; Nagendra *et al*. 2014). Land clearing for agriculture and urban expansion, for instance, have resulted in fragmentation of forest biomes across the world (Reinmann *et al*. 2020). This practice also leads to increased pollutants, like nitrate and ammonium, which in turn, change soil biogeochemical properties and microbiome function (Wang *et al*. 2017). Additionally, industrial activities can lead to heavy metal and organic contamination in urban soils, in particular polycyclic aromatic hydrocarbons (PAHs), that are toxic to humans and are added to soil through processes like burning fossil fuels and waste incineration (Fernández-Luqueño *et al*. 2011; Delegan *et al*. 2022). Finally, urban land use and management practices like fertilization, irrigation, and the installation of impervious surfaces increase nutrient inputs into soils via runoff (Fenn *et al*. 2003). Nitrogen runoff in particular can enhance microbial denitrification processes that release nitrous oxide – a potent greenhouse gas (Kettering *et al*. 2013; Toor *et al*. 2017).

In parallel, soil microorganisms (i.e., bacteria, archaea, and fungi) are major conductors of global carbon and nutrient cycles. Soil biogeochemical properties that are impacted by urbanization like moisture, temperature, pH, and carbon and nutrient concentrations have a strong effect on microbial communities (Rousk *et al*. 2010; Brockett, Prescott and Grayston 2012; Karhu *et al*. 2014; Graham *et al*. 2016; Widdig *et al*. 2020). Microbial diversity is also dependent on interspecies and plant-microbe interactions (Wilpiszeski *et al*. 2019; Sharma *et al*. 2020; McClure *et al*. 2022), which have been shown to increase the relative abundance of genes involved in greenhouse gas emissions in urban soils (Delgado-Baquerizo, Eldridge and Yu-Rong 2021). More generally, urban soil microbiomes have been shown to exhibit homogenization of bacterial communities compared to surrounding natural ecosystems (Gill *et al*. 2020; Delgado-Baquerizo, Eldridge and Yu-Rong 2021).

The extent to which the urban environment drives soil microbiomes towards a common state (i.e., microbiome composition and function; urban convergence hypothesis) versus supporting a robust, diverse microbial community (i.e., the urban mosaic) remains debated (Pouyat *et al*. 2015). Within the dataset re-analyzed here, for example, it was reported that soil biogeochemical properties like organic carbon and total nitrogen content converged due to anthropogenic factors (Pouyat *et al*. 2015). Other studies have also seen the effect of urban convergence on soil properties, such as soil particle size and soil carbon content (Herrmann, Schifman and Shuster 2020). However, due to the plethora of land covers and uses contained within different metropolitan areas, it remains to be explored whether or not urbanization has a uniform effect that will drive microbial communities towards greater biodiversity loss or greater taxonomic and functional potential. Further, understanding the extent to which urbanization acts upon microbial taxa versus the functional traits they contain is important for representing the impacts of population growth on global biogeochemistry. Standardized spatial observations distributed within and across cities are key to resolving these questions.

Determining a core soil urban microbiome is essential to study patterns associated with anthropogenic factors. Broadly defined, a core microbiome is a set of shared taxa and/or functional potential across all samples in a dataset or a group within a dataset (Turnbaugh *et al*. 2007; Hamady and Knight 2009; Shade and Handelsman 2012; Neu, Allen and Roy 2024). Microorganisms consistently found in all assemblages within a specific habitat have been proposed as essential for the community’s functioning (Turnbaugh and Gordon 2009; Vael and Desager 2009; Benson *et al*. 2010; Sommer and Bäckhed 2013). Therefore, pinpointing specific taxa and functional potential in a core microbiome is the initial step in identifying a ‘healthy’ community and forecasting how the community will react to disturbances. Methods to determine a core microbiome can vary with experimental factors such as spatial or temporal scale and can leverage occurrence-based data, relative abundance data, or a combination of relative abundance and occurrence data (Shade and Handelsman 2012; Neu, Allen and Roy 2024). Here, we define the core urban soil microbiome as organisms and functional genes that are distributed across global urban land uses and are also scarce or absent from more pristine reference soils.

We use metagenomic sequences from soils collected by the Global Urban Soil Environment Ecology Network (GLUSEEN; Pouyat et al., 2017) to investigate the taxonomic and functional diversity of the urban soil microbiome. We aim to examine if soil taxonomic and functional diversity are conserved across worldwide urban cities, if changes in taxonomy and functional potential are coupled, and if there are ubiquitous (“core”) taxa and functions that are characteristic of urban soils regardless of climate and geography. Previous work with this dataset, by Epp Schmidt, et al. (2019, 2017) has shown some consistency in archaeal and bacterial communities, with archaeal convergence being driven by ammonia- and methane-oxidizing organisms and genes and bacterial convergence being driven by nitrite-oxidizing organisms. However, Epp Schmidt, et al. (2019, 2017) did not explore aspects of urban microbiomes related to contaminants and other carbon and nitrogen cycling processes. We further extend this work by reporting concomitant changes in taxonomy and functional diversity within and across cities; extracting the core urban microbiome across all cities; and uncovering a newly reported set of widespread soil genomic potential for the tolerance and/or cycling of urban pollutants.

## Methods

### Dataset Description

We used publicly available metagenomic sequencing data from the Global Urban Soil Environment Ecology Network (GLUSEEN) to investigate the urban soil microbiome from 66 samples across five global cities: USA: Baltimore (n=11), Finland: Helsinki (n=12), Finland: Lahti (n=14), Hungary: Budapest (n=14), and South Africa: Potchefstroom (n =15) (Figure 1). Soils spanned three levels of urbanization that encompass a gradient of common anthropogenic impacts: (1) remnant: within city limits, (2) turf: managed land, and (3) ruderal: high degree of physical disturbance; in addition to reference soils with low urban impacts near each city. Further details on land use classifications, soil collection, and metagenomic sequencing are described by Pouyat, et al. (2015) and Epp Schmidt, et al. (2017). Briefly, soils were collected prior to 2017 from 0-10 cm depth, and shotgun metagenomic sequencing was performed on Illumina HiSeq 3000 using standardized methods. Samples used for this publication are available at MG-RAST (Yarwood and Epp Schmidt). (https://www.mg-rast.org/mgmain.html?mgpage=project&project=mgp16346).

**Figure 1.**
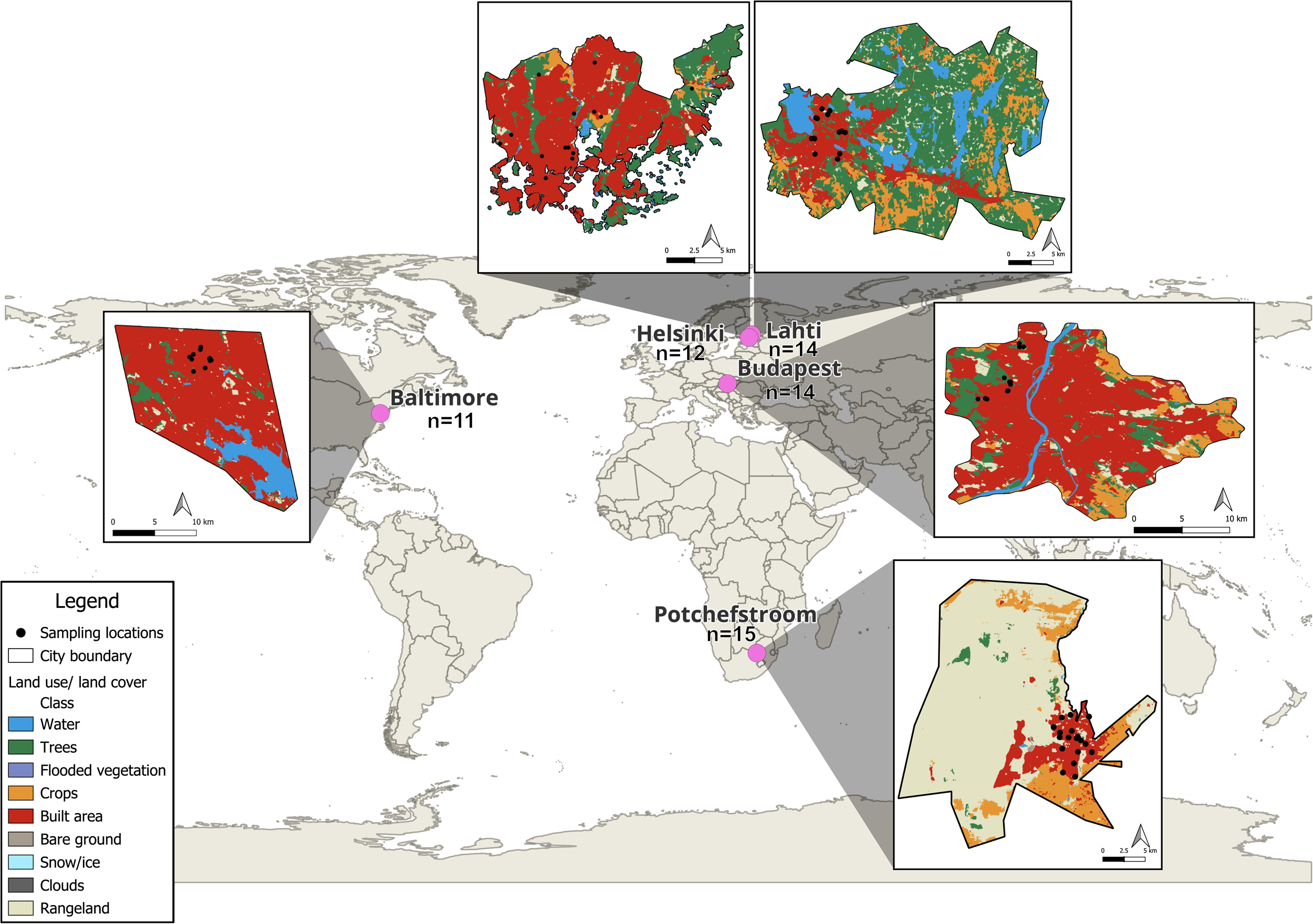
Geographic distribution and land use/land cover of urban soil microbiomes. All soils are from locations with a built area land use/land cover classification according to ESRI (Kontgis et al., 2017; 2021). Cities are colored by land use/land cover categories, and sampling locations are denoted by black dots. Land use/land cover distributions for each city are derived from Environmental Systems Research Institute, ESRI (2017 data; Kontgis et al., 2017; 2021). Land use analyses were performed in QGIS 3.32.3-Lima equipped with a semi-automatic classification plug-in.

### Taxonomic and Functional Inference of Metagenomes

Metagenomic sequences from each soil were retrieved from Metagenomic Rapid Annotations using Subsystems Technology (MG-RAST) and processed using the Joint Genome Institute (JGI) Integrated Microbial Genomes and Microbiomes (IMG/M) standard workflows (Clum *et al*. 2021). Briefly, reads meeting the QC standards were assembled into contigs using BBDuk version 38.79, bbcms version 38.44 and metaSPADES version 3.13.0 (Nurk *et al*. 2017; Bushnell 2023). Read-based taxonomy was assigned to assembled contigs using the genome-based taxonomy database, and assembled contigs were further annotated against the Kyoto Encyclopedia of Genes and Genomes (i.e., KEGG Orthology, KO; (Kanehisa et al., 2019, 2000, 2023)) as part of standard JGI workflows (Parks *et al*. 2018; Chaumeil *et al*. 2020). While read-based taxonomy is a useful tool for determining microbial clades via metagenomics, future work that assesses taxonomy by sequencing selected portions of conserved marker genes may be a useful complement to the results presented here (Wood and Salzberg 2014; Wood, Lu and Langmead 2019; Clum *et al*. 2021). After sequence processing, we summarized taxa and KO for each soil sample into matrices by counting sequences in each soil assigned to every unique taxon (order-level due inconsistencies in taxonomic annotations extending beyond microbial orders) or KO. Counts in each matrix were normalized using a relative log expression (RLE; Pereira et al., 2018) implemented in the R package ‘DESeq2’ (Love, Huber and Anders 2014). KO were mapped to pathways using the ‘KEGGREST’ package in R (version 4.3.1) (Tenenbaum and Maintainer 2023). Code is available on Figshare (https://figshare.com/s/d788b28a869481c25689).

### Statistical Analysis

All statistical analyses were implemented in R (version 4.3.1) with base statistical packages and with the ‘vegan’ and ‘usedist’ packages (Bittinger 2020; Oksanen *et al*. 2022). We examined how taxonomic diversity varied across land uses and cities by calculating alpha-(richness and Shannon Diversity Index) and beta- diversity (Bray-curtis dissimilarity), followed by ANOVA and Tukey’s honestly significant difference (HSD) post hoc test or by permutational multivariate analysis of variance (PERMANOVA) to test for statistically significant differences in alpha- and beta-diversity, respectively. Diversity metrics were independently calculated based on the distribution of microbial orders (for microbial taxonomy) and KO (for microbial functional potential). Then, we individually compared how taxonomic and functional composition changed with geographic distance, using the Haversine dissimilarity matrix within and across cities, using Mantel tests. Relative changes in taxonomic and functional composition were assessed using Mantel test of Bray Curtis dissimilarity for microbial taxonomy vs. functional potential.

Additionally, we visualized data in the following ways. For visualizations of read-based taxonomy, we first generated phylogenetic trees using phyloT v2 (phyloT: a phylogenetic tree generator), an online tree generator based on the Genome Taxonomy Database. Then, we visualized the tree in R using the packages, ‘ggtree’ (Yu *et al*. 2017), ‘treeio’ (Wang *et al*. 2020), and ‘ggnewscale’ (Campitelli 2020). We generated heatmaps using the ‘pheatmap’ package (Kolde 2019). Distributions of shared and unique taxa and KO across cities were visualized using the ‘ComplexUpset’ package in R (Krassowski 2020; Lex *et al*. 2020). All other visualizations were generated with ‘ggplot2’ (Wickham 2016).

## Results

### Microbial Diversity of Urban Soils

When comparing diversity metrics across cities, the Baltimore soil microbiome appeared to be distinct. The taxonomic diversity of soil microbiomes in Baltimore was both higher (alpha) and distinct (beta) compared to other cities (Figure 2). Excluding reference soils, taxonomic richness was nearly two-fold higher in Baltimore soils than all other cities (261.46, vs. 120.57-172.20, P < 0.001) and Shannon diversity followed a similar trend (**H’**=3.77, vs. **H’**=0.80-1.87, P < 0.001, Figure 2). Baltimore soils also had lower functional richness compared to Budapest and Potchefstroom (mean = 2,189.636 vs. 3440.714 and 3325.067 unique KO per sample, P= 0.021 and 0.040, respectively) and the lowest Shannon diversity index when compared to all other metropolitan areas (**H’**= 7.110, **H’** range= 7.669-7.816, P<0, Figure 2). Taxonomic and functional composition in the Baltimore soil microbiome was also distinct when compared with other cities (PERMANOVA, both with and without reference soils P < 0.001, Figure 2).

**Figure 2.**
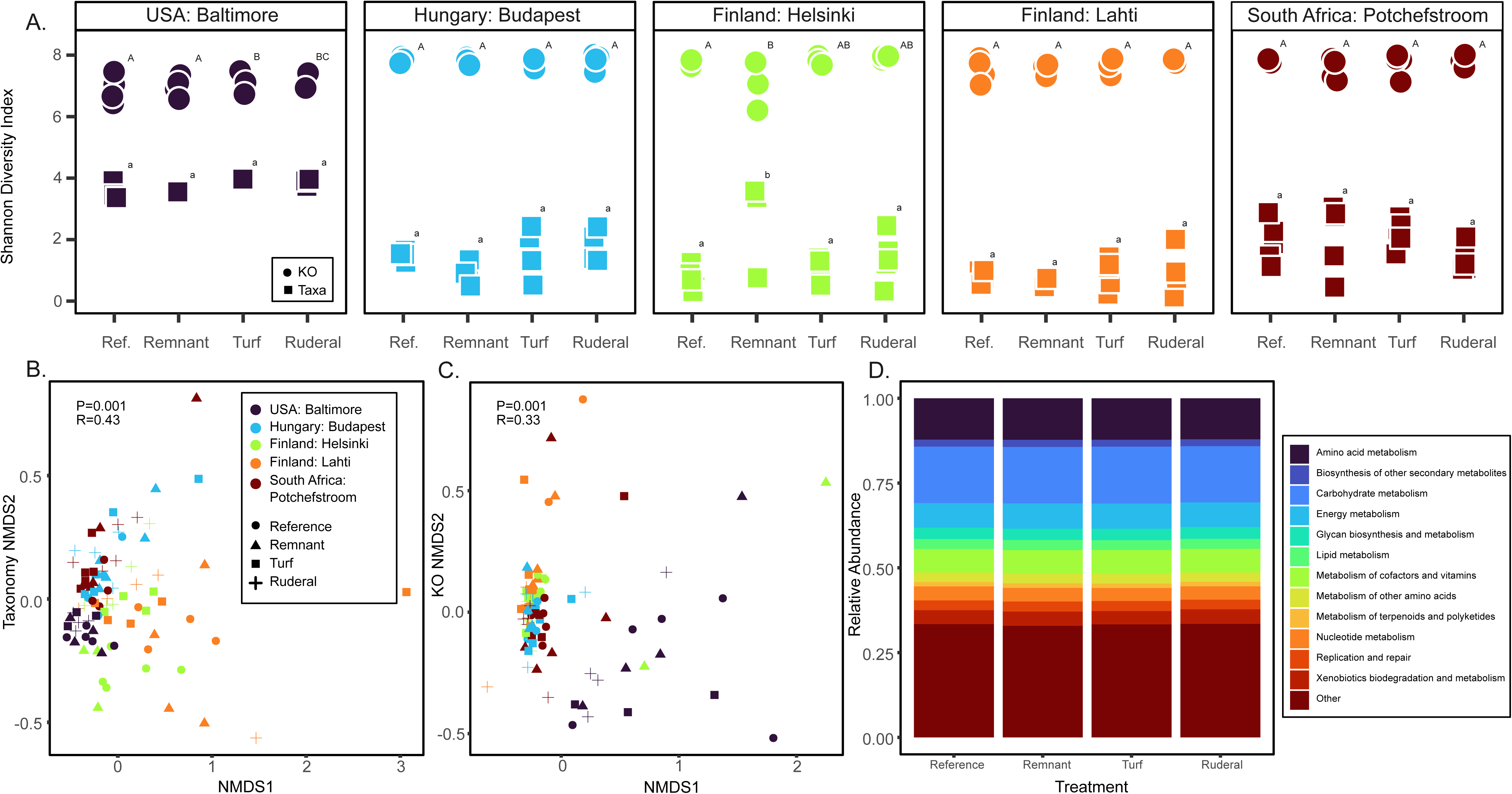
Alpha and beta diversity of microbial taxonomy and KO. Shannon Diversity Index (A) and Non-metric MultiDimensional Scaling (NMDS) for each sampling location for taxonomic order (B) and KO (C), colored by city and with distinct shapes for treatment. The top 10 most abundant (D) metabolic pathways are shown with stacked bar plots. In (A), letters represent statistically distinct groupings (P=<0.05). In (B) geographic location accounts for 43% of the variation seen for taxonomic order and 33% of the variation seen for KO (P=0.001, respectively).

Highly urbanized land uses in most cities contained distinct soil microbiomes relative to reference soils. Though there was no significant change in taxonomic or functional alpha diversity metrics within urban land uses (ruderal, remnant, turf) and only slight differences in alpha diversity between urban and reference land uses (Figure 2); soil microbiomes associated with the most highly urbanized land use (ruderal) were distinct from reference soils with respect to taxonomy and/or functional potential in Baltimore, Potchefstroom, and Helsinki (PERMANOVA, P = 0.02, 0.02, and 0.09 for taxonomy and P = 0.02, 0.18, and 0.01 for functional potential, respectively).

We then tested relationships between microbial diversity (taxonomic and functional) and geographic position, and between taxonomic and functional dissimilarity, to assess the extent to which taxonomic and functional attributes of urban microbial communities were coordinated through space (Figure 3). When examining soils from all cities collectively, both taxonomic and functional dissimilarity increased with geographic distance (P= 0.01), an effect that was present both with and without the inclusion of reference soils. For each city individually, neither taxonomic nor functional diversity varied significantly with geographic distance within urban land uses in any city (P > 0.10). We obtained similar results when including reference soils, where the only significant correlations with distance were found in Baltimore (P = 0.04, 0.05 for taxonomy and functional potential, respectively). However, taxonomic and functional composition were partially coupled across all cities collectively (all P = 0.01, with and without reference soils) and within Budapest, Helsinki, and Potchefstroom (all P=0.01, with and without reference soils).

**Figure 3.**
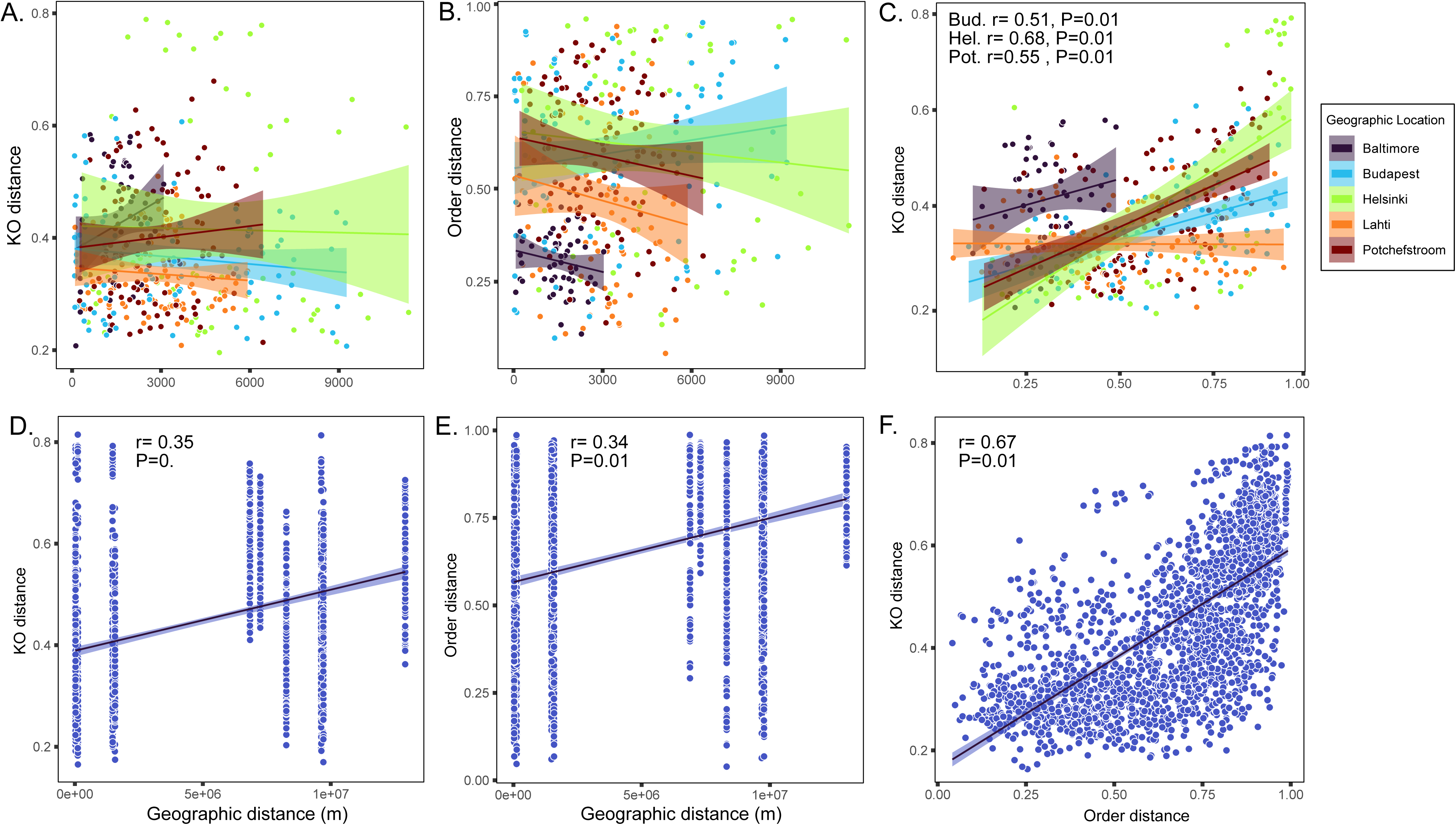
Correlations between urban soil microbiome taxonomy, functional potential, and geographic distance. (A-C) show correlations within individual cities, denoted by color. (A) and (B) show dissimilarity in the composition of KO’s and microbial order, respectively, vs. geographic distance. (C) shows KO vs. taxonomic dissimilarity. (D) and (E) show dissimilarity in the composition of KO’s and microbial order, respectively, vs. geographic distance across all cities. For all plots, geographic dissimilarity is calculated using Haversine distances, and KO and taxonomic dissimilarity are calculated with Bray-Curtis dissimilarity. Statistical significance was determined using Mantel tests. Taxonomy is grouped at the order-level for all analyses.

### Core Urban Microbial Taxa and Metabolic Potential

Two hundred fifty-seven microbial orders were common to soil microbiomes across all cities (hereafter, ‘core’ organisms, Figure 4). Although most of these orders were also contained within reference soils, they exhibited clear increases in abundance from reference to more urbanized soils (Figure 5, Figure S1). Among core urban microbial orders, eight bacterial and two archaeal orders had greater than 50% of their total abundance contained within ruderal soils (the most urban land use type, unclassified Candidatus Saccharibacteria, unclassified Crenarchaeota, Caldisericales, Clostridiales, Desulfurellales, Petrotogales, Micrococcales, Archaeoglobales, Chlamydiales, and unclassified Firmicutes), with low abundance in reference soils (0-15% for eight orders and 17-29% for the remaining two orders; Table S1). Eleven more orders (incl. seven archaeal orders) had more than 50% of their total abundance contained within turf soils, with low corresponding abundance in reference soils (<12% of total abundance contained within reference soils, Aeromonadales. Caulobacterales, Burkholderiales, Candidatus Nitrosocaldales, Methanopyrales, unclassified Thermoplasmata, Nitrosopumilales, unclassified Aigarchaeota, unclassified Calescamantes, Thermoplasmatales, unclassified Thaumarchaeota). Overall (regardless of presence in reference soils), the most abundant shared orders of bacteria included Enterobacterales, Burkholderiales, Lactobacillales, Bacillales, Rhizobiales, Micrococcales, Caulobacterales, Acidobacteriales, Clostridiales, and Pseudomonadales.

**Figure 4.**
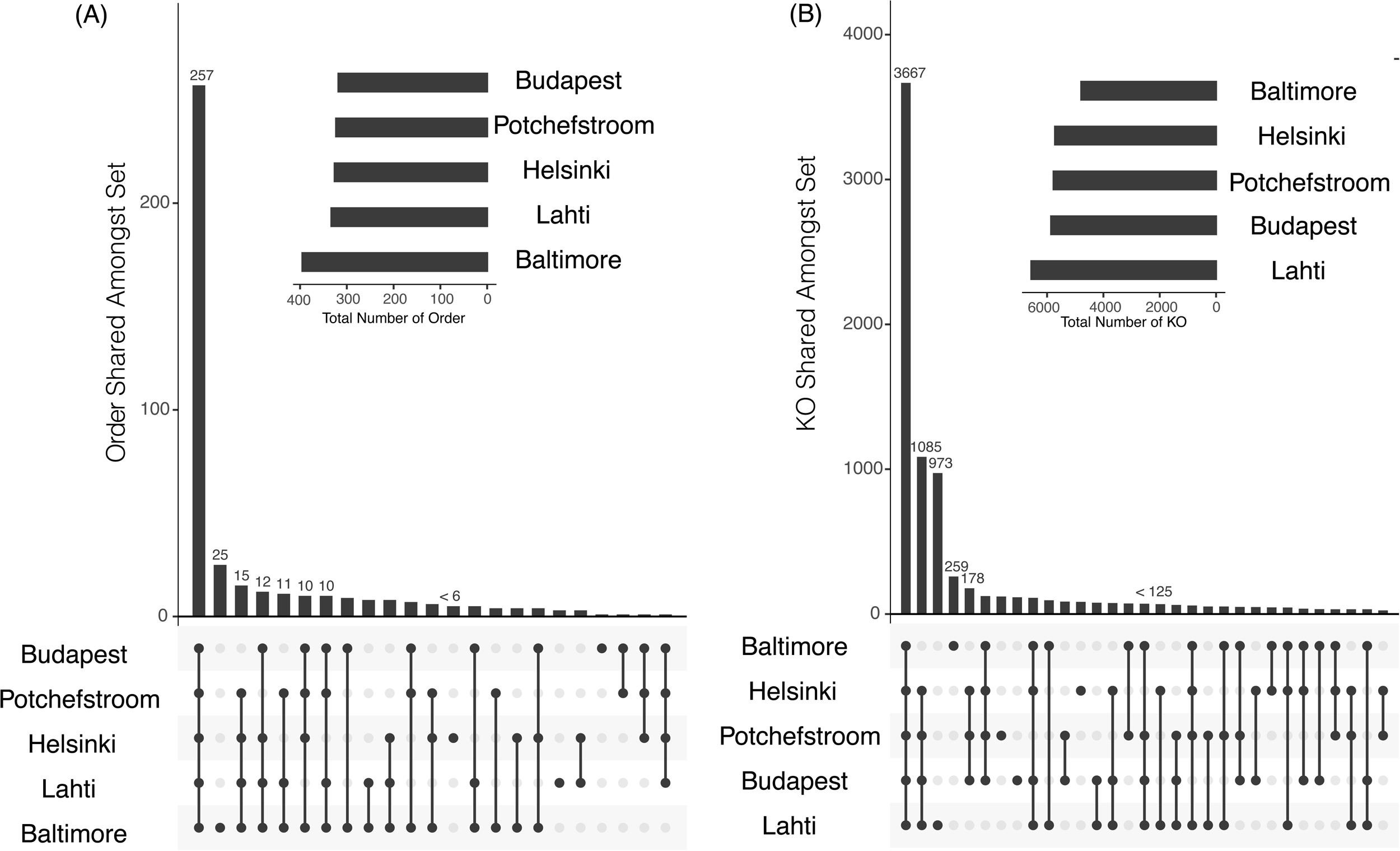
Microbial organisms and functional potential across cities. (A) and (B) show UpSet plots of microbial taxonomy (order-level) and functional potential (KO) respectively. Sets of intersections between cities are shown by dots on the x-axis. The number of shared (A) microbial orders or (B) KO for each intersection is denoted by bar height. Insets show the total number of microbial orders or KO per city.

**Figure 5.**
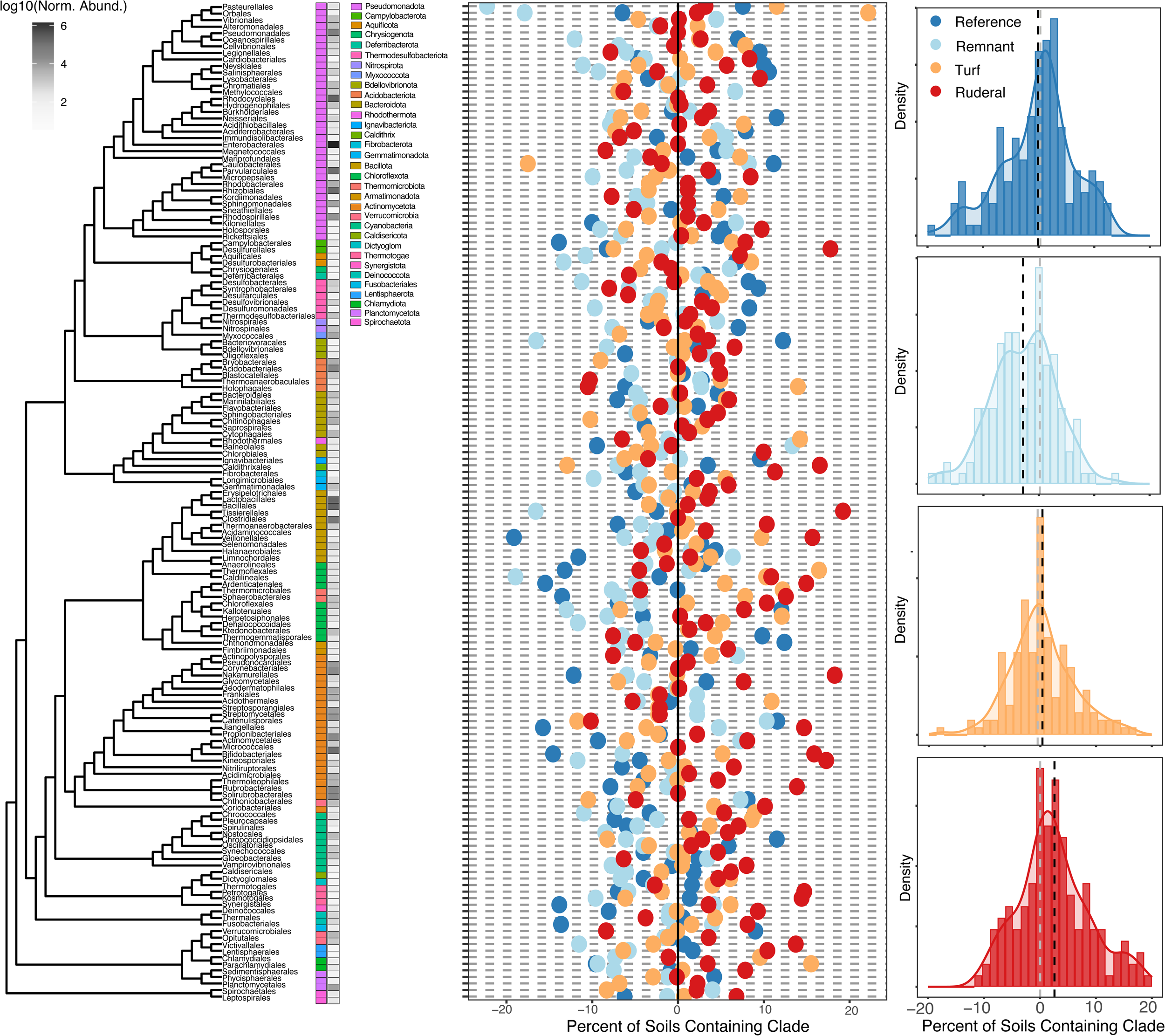
Ubiquitous urban soil bacteria. (A) Phylogenetic tree of soil bacteria found across all five cities. Color bars denote phyla. Grayscale heatmap shows the normalized abundance of each bacterial order in urban soils (i.e., all soils except reference soils). (B) Cleveland dot plot showing the percent of soils containing each microbial order in each soil type. The mean across all samples is set at zero (black line) and over/underrepresentation is denoted by dots further away from the center line. Land use type is depicted by colors, warmer colors indicate more urban land cover. (C) Histograms showing the distribution of data in (B) for each land cover type. The vertical gray line represents zero deviation from the mean of all samples (equivalent to the black line in C). The mean of each land cover type is shown with a dashed vertical line of that color. Histograms with means to the right of 0 suggest an overrepresentation of selected bacterial orders in the associated land cover type.

**Figure 6.**
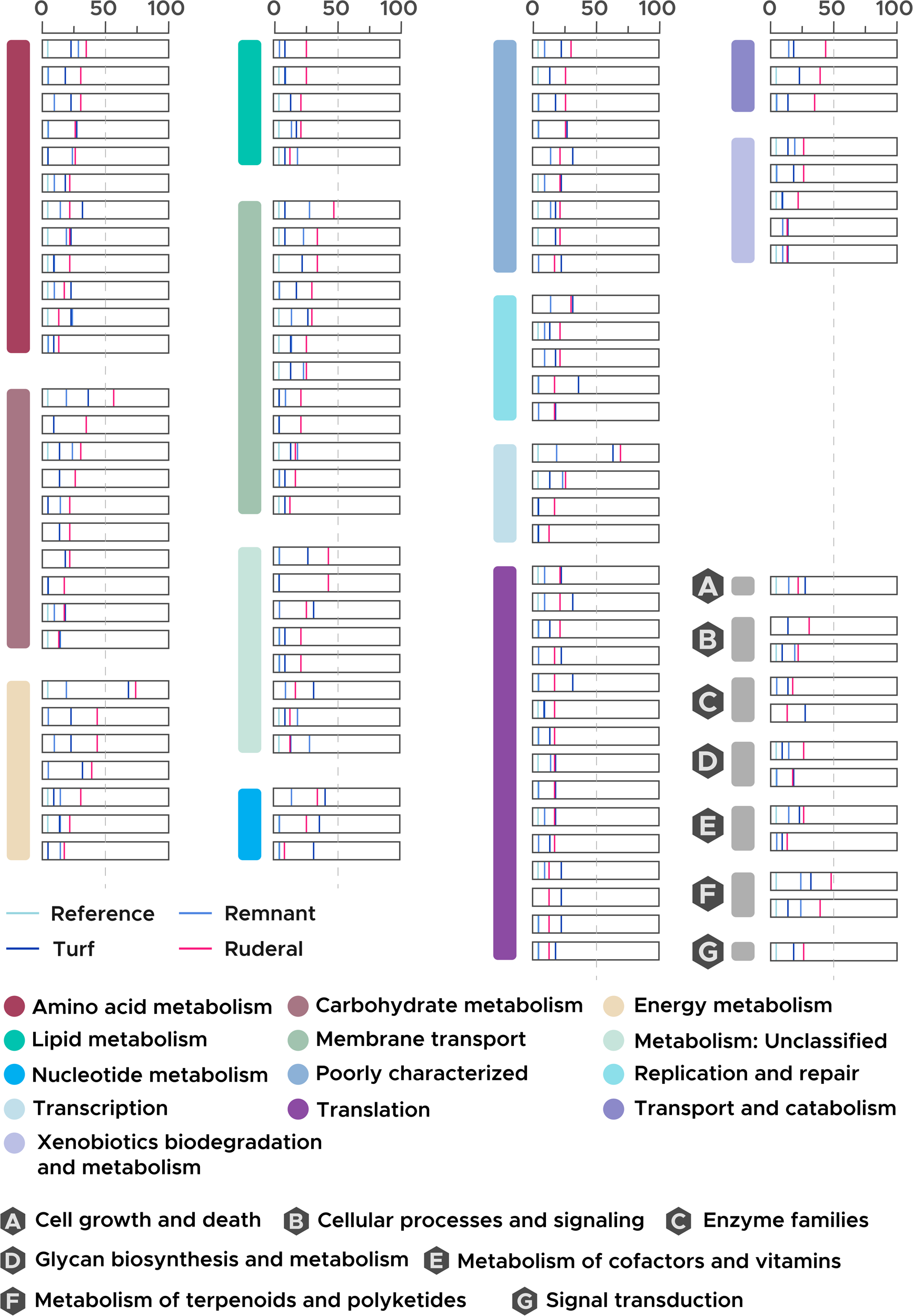
Selected metabolic pathways of urban soil microbiomes that are found in less than 5% of reference soils. Selected pathways were manually reconstructed from KEGG pathways and are color-coded as indicated in the legend.

Core urban archaeal orders were generally more evenly distributed across reference and urban land uses, however, several taxa still appeared to be more prevalent in urbanized soils. No core orders were found only in urban land uses, but 7 of 18 core archaeal orders were at least twice as abundant in ruderal vs. reference soils: Methanopyrales, Nitrosopumilales, Nitrososphaerales, Thermoproteales, Thermoplasmatales, Archaeoglobales, and Methanocellales. Overall (regardless of presence in reference soils), Nitrososphaerales was by far the most abundant shared archaeal order, followed by other nitrogen- and methane-cycling (Nitrosopumilales, Methanosarcinales, Methanomicrobiales, Methanobacteriales) and salt-tolerant archaea (Halobacteriales, Haloferacales, Natrialbales). Interestingly, after excluding reference soils, Baltimore soils displayed five times as many auxiliary (i.e., not in core microbiome) microbial orders as any other city (25 vs. five or less in all other cities; Figure 4).

Further, we found 3,667 KO shared across urban land uses in all cities (Figure 4). Regardless of their presence in reference soils, the top 10 most abundant core KO encoded proteins that are associated with generalized cell functions (K07347 outer membrane usher protein, K01992, K02004, and K01990 ABC-2 type transport system permease and ATP-binding proteins, K02529 LacI family transcriptional regulator, galactose operon repressor, K03088 polymerase sigma-70 factor ECF subfamily, K01652 acetolactate synthase I/II/III large subunit [EC:2.2.1.6], K07497 putative transposase), notably including two metal-dependent functions: K02014 (iron complex outer membrane receptor protein) and K02035 (peptide/nickel transport system substrate-binding protein).

When narrowing these potential functions to those not widely contained within reference soils, our results point to a core set of 111 gene annotations that were found in urban land uses in all five cities and in less than 5% of reference soils. In particular, 29 KO were common to urbanized land in all cities but were not found in any reference soil. Of these, 12 were found in at least 25% of soils from the most urbanized land use type (ruderal). This included 4 genes associated with nitrogen cycling (K02591 *nifK*, nitrogenase molybdenum-iron protein beta chain, K02586 *nifD*, nitrogenase molybdenum-iron protein alpha chain, K02588 *nifH*, nitrogenase iron protein NifH, K12256 *spuC* putrescine---pyruvate transaminase [EC:2.6.1.113]). In addition to the two Mo-dependent genes related to N cycling (*nifK* and *nifD*), five other genes in this group are closely associated with heavy metals, contaminants, and/or microbial stress responses (K05297, rubredoxin---NAD+ reductase; K18601 aldehyde dehydrogenase [EC:1.2.1.-]; K07231 putative iron-regulated protein, K03332, *fruA*, fructan beta-fructosidase [EC:3.2.1.80], K01960, *pycB* pyruvate carboxylase subunit B). The three remaining genes were associated with common cellular functions (K19123 *casA*, CRISPR system Cascade subunit CasA, K03041, *rpoA1*, DNA-directed RNA polymerase subunit A’ [EC:2.7.7.6], K10726, replicative DNA helicase Mcm [EC:5.6.2.3]).

If we further extend this analysis to urban soil KO found in less than 5% of reference samples, we generate an additional 82 KO for consideration, of which 34 are found in more than 25% of soils from the most urbanized land use type (Table S1). Notable KO include: K08352 (thiosulfate reductase/polysulfide reductase chain A [EC:1.8.5.5], in 74% of ruderal soils); K13794 (LysR family transcriptional regulator, regulatory protein for *tcuABC*, in 70% of ruderal soils); K00869 and K00938 (mevalonate kinase [EC:2.7.1.36] and phosphomevalonate kinase [EC:2.7.4.2], in 48% and 39%); K02587 and K02229 (*nifE*, nitrogenase molybdenum-cofactor synthesis protein NifE, and *cobG*, precorrin-3B synthase [EC:1.14.13.83], in 44% and 26%); K00520 and K08365 (mercuric reductase [EC:1.16.1.1] and MerR family transcriptional regulator, mercuric resistance operon regulatory protein, in 44% and 26%); K18833 (MFS transporter, DHA3 family, multidrug efflux protein, in 35%); K09819 (manganese/iron transport system permease protein, in 26%); K00596 (2,2-dialkylglycine decarboxylase (pyruvate), in 30%); genes related to fructose/fructan (K10554, *frcA* fructose transport system ATP-binding protein and K00895 diphosphate-dependent phosphofructokinase [EC:2.7.1.90], in 30% and 30%), K16876 (2-furoate---CoA ligase [EC:6.2.1.31], in 26%); genes related to glycerol contamination (K17325 glycerol transport system ATP-binding protein in 30%, K00712 *tagE* poly(glycerol-phosphate) alpha-glucosyltransferase [EC:2.4.1.52] in 26%, K17322 *glpP* glycerol transport system permease protein in 26%), signatures of viral infection (K01186, sialidase-1 [EC:3.2.1.18] in 26%, K19046, *casB*, CRISPR system Cascade subunit CasB, in 39%), some uncharacterized functions, and several functions related to amino acid transport and metabolism.

Despite having the fewest auxiliary functional genes, Baltimore soils harbored a distinctly unique microbiome enriched in genes associated with heavy metal resistance and toxin processing, in contrast to other cities where auxiliary functions were dominated by common cellular processes. Excluding reference soils, Lahti soils had the most auxiliary microbial KO at 2895, followed by Budapest (2190), Helsinki (2104), Potchefstroom (2053), and Baltimore (1122). While the auxiliary microbiome of Baltimore soils contained the fewest potential functions of any city, it was also mostly unique in comparison to other cities. The most abundant of these auxiliary potential functions is thought to convey heavy metal resistance (K15725, outer membrane protein, heavy metal efflux system). Other highly abundant auxiliary functions in Baltimore soils were also associated with metals or other toxins, such as: K01578 MLYCD, malonyl-CoA decarboxylase [EC:4.1.1.9], K15855, exo-1,4-beta-D-glucosaminidase [EC:3.2.1.165], K01577 *oxc*, oxalyl-CoA decarboxylase [EC:4.1.1.8], K04108 4-hydroxybenzoyl-CoA reductase subunit gamma [EC:1.1.7.1],and K01301 N-acetylated-alpha-linked acidic dipeptidase [EC:3.4.17.21]. In contrast, the auxiliary microbiome of other cities was mostly comprised of common cellular processes (K01223, K02761, K07345, K02073, K03475).

## Discussion

Urban centers are increasing in extent and population density, in turn disturbing soil microbiomes and the biogeochemical processes that they mediate (Christel *et al*. 2023). Soil microbial functions are an essential component of global biogeochemical cycles, as microorganisms regulate soil C storage and respiration (Gougoulias, Clark and Shaw 2014). Here, we used 89 soil metagenomes from the GLUSEEN network to evaluate patterns in the taxonomic and functional potential of urban soil microbiomes. We extend the work of Epp Schmidt by revealing a suite of core urban microorganisms and metabolisms related to (1) trace and heavy metals, (2) fructose/fructan, (3) viral infection, and (4) glycerol; in addition to N cycling processes that have been reported previously. These novel findings present a basis for understanding and modeling the ways in which urban biogeochemical cycles are distinct from natural ecosystems and for sustainably managing growing population centers.

### Taxonomy and Functional Potential of Urban Soil Microbiomes

We found biogeographical patterns in the urban soil microbiome that align with ecological processes such as microbial niche diversification (urban mosaic hypothesis) and environmental filtering (urban convergence hypothesis). There was a great array of functional potential relative to the amount of taxonomic diversity present in the urban soil microbiome (Figure 2). High functional diversity lends support for the urban mosaic hypothesis, which proposes that microbial diversity increases with urbanization as a result of interacting selective pressures like land use, nutrients, pollution, and other factors on microbiomes (Christel *et al*. 2023; Scholier *et al*. 2023). Under the urban mosaic hypotheses, potential microbial functions, and in some cases taxonomic composition, expand to occupy new niche space opened by anthropogenic activity. For example, Mo, et al. and Scholier, et al. (2024; 2023) reported increased soil microbiome alpha diversity in anthropogenic areas compared to natural areas. Taxonomic similarities, on the other hand, have been more widely reported across cities (McKinney 2006; Luck and Smallbone 2010; Danko *et al*. 2021; Lokatis and Jeschke 2022). This provides evidence towards the urban convergence hypothesis, where physical characteristics of the urban environment drive microbial similarities, and supports the existence of a core urban microbiome. In our study, we observed comparatively low taxonomic diversity in urban soils (Figure 2) and a clustering of microbiomes from highly urbanized land uses across cities (Figure 2, NMDS) that lends support for common microbial taxa and functional potential across five vastly different cities. This is consistent with findings in urban greenspaces where despite greater functional diversity, microbial taxonomies are relatively homogenized (Delgado-Baquerizo, Eldridge and Yu-Rong 2021; Mo *et al*. 2024).

Interestingly, Baltimore appeared to have a distinct soil microbiome relative to all other urban centers. It had the highest taxonomic diversity but also the least functional diversity as compared to other cities (which had similar diversity, Figure 2). Baltimore was also the only city that showed a sizable increase in taxonomic diversity in more urban (turf and ruderal) vs. more pristine (reference and remnant) soils (Figure 2). We posit that the influence of environmental factors on soil microbiomes may be particularly important in Baltimore, which has a long history of industrialization, leaky sewage infrastructure, and high pollution levels (Yesilonis, Pouyat and Neerchal 2008; Dickerson *et al*. 2024; *Baltimore’s Wastewater Treatment Plants*). For example, Baltimore has been shown to display a dramatic urban heat island effect with increased temperatures along an urban gradient (Corpuz *et al*. 2024) and has reportedly high levels of lead and other pollutants (Yesilonis, Pouyat and Neerchal 2008; Schwarz, Pouyat and Yesilonis 2016). Each of these factors may both exert unique selective pressures on microbial functions and vary substantially at the local scale. They may also lead to the emergence of genetic potential that is not well-described in microbial communities (i.e., microbial dark matter), which would result in the suppression of detectable functional potential observed in this study. While the reasons driving different patterns in microbial taxonomic and functional diversity in Baltimore are unknown, its distinctive patterns suggest further investigation into the unique taxonomic and functional potential its soil microbiomes contain.

### Spatial variation in functional potential and taxonomic composition across the urban soil microbiome

While the mixture of diverse land uses within cities may obscure patterns based on space alone, the low magnitude of compositional changes within and across cities further underscores the existence of a core urban microbiome. Differences between urban and reference soils suggest that these core organisms and functions are distinct from more natural environments (Figure 3). We found that urban soil microbiome composition and functional diversity tended to be consistently structured through space, consistent with previous work on this dataset (Epp Schmidt *et al*. 2019). For each individual city, neither taxonomic nor functional diversity varied significantly across spatial distances. Across cities, both taxonomic and functional dissimilarity showed some correlation with spatial distances, 33.7% and 34.6% respectively (Figure 3). This variation in microbial community composition across cities was likely due to overarching drivers such as soil mineralogy, pH, weather, and other abiotic factors that are known to influence soil microbiomes (Brockett, Prescott and Grayston 2012; Cao *et al*. 2016; Wang *et al*. 2019; Widdig *et al*. 2020).

Interestingly, microbial taxonomic composition and functional potential were partially decoupled from each other and showed different biogeographical relationships (Figure 3). Taxonomic composition was more variable than functional potential through space, both within and across cities (as indicated by flatter slopes on Figure 3), and was virtually uncorrelated to functional potential in Lahti and Baltimore. Microbial functional potential is expected to be somewhat correlated to taxonomic composition due to phylogenetic conservation of microbial traits, which can lead to similar responses to environmental factors like substrate availability and climatic variables (Martiny, Treseder and Pusch 2013). However, if ecological selection operates on microbial genomic elements that are exchanged between organisms or if selection operates on unrelated genes and organisms, shifts in microbial taxonomy may not be coordinated with changes in genomic potential (Knelman and Nemergut 2014; Bier *et al*. 2015; Graham *et al*. 2016; Graham and Knelman 2023). We posit that the diversity of factors such as available substrates, carbon sources, and/or pollutants in the urban environment may lead to variation in gene presence and abundance despite taxonomic convergence. Given the usage of microbial taxa as biomarkers of ecosystem health (Dubey *et al*. 2019) and as engineering solutions (Bala *et al*. 2022), the disconnect between taxonomy and potential function observed here for urban soils is an important consideration in sustainable management practices.

### Ubiquitous (core) vs. auxiliary taxa and metabolisms in global cities

Examining the core urban microbiome provides insight into the microorganisms that are likely to mediate the biogeochemical cycling of key nutrients, greenhouse gasses, and pollutants in urban soils. Bacterial orders that were ubiquitous across cities and urban land covers relative to reference soils were typified by microorganisms usually associated with plant health, rhizosphere functions, and antibiotic and/or contaminant production and consumption (Figure 5, Figure S1). For example, Enterobacterales and Rhizobiales reside in the root systems of plants and some members have been found to maintain plant health under abiotic stress (Garrido-Oter *et al*. 2018; Andrés-Barrao *et al*. 2021). *Bacillus*, a common microorganism in the Bacillales order, is also ubiquitous in soils and promotes plant health through nutrient cycling and pathogen protection (Garbeva, van Veen and van Elsas 2003; Liu *et al*. 2019). With regards to antibiotic and contaminant metabolism, some members in the Enterobacterales order (e.g., *Enterobacteria*) harbor increased antibiotic resistance genes in urban centers, while others are human pathogens (e.g., *Salmonella*, *Escherichia*, *Shigella*). Burkholderiales have also been shown to participate in pentachlorophenol degradation, a chemical used as a pesticide that persists in soil and has been coupled to nitrogen, sulfate, and iron reduction (Tong *et al*. 2015; Kang *et al*. 2018; Fresia *et al*. 2019); while Clostridiales and Caulobacterales are commonly associated with a wide range of contaminants (Abbasian *et al*. 2016; Thomas and Cébron 2016). Like bacteria, ubiquitous urban archaea have been previously associated with nitrogen and methane cycling and/or salt tolerance (Fenn *et al*. 2003; Wang *et al*. 2013; Asabere *et al*. 2018; Strokal *et al*. 2021). While we cannot directly assess the coupling of these urban organisms with trace metals, pollutants, and/or harsh conditions using the data available in this study, their high plasticity and potential to acquire new genes (Brunder and Karch 2000; Justice *et al*. 2008; Liu *et al*. 2019) may make them a key focus for future research.

We also report core soil microorganisms with greater than 50% of their abundance contained in ruderal soils and corresponding low abundance in reference soils as organisms for deeper investigation (Figure 5, Figure S1). We focus on ruderal soils because they represent the most urbanized land cover type investigated here. First, Saccharibacteria has been reported to be a key member isolated from activated sludge (Ferrari *et al*. 2014; Kindaichi *et al*. 2016) and is often found in soil and soil-like environments. Members of Saccharibacteria are able to metabolize diverse compounds, putatively conveying an ability to thrive in a polluted urban environment (Kindaichi *et al*. 2016). Saccharibacteria are a common constituent of the human oral microbiome which may also explain its presence in urban environments that have high human populations (Bor *et al*. 2019). Members of Clostridiales and other Firmicutes are also commonly associated with contaminant degradation, including polycyclic aromatic hydrocarbons (PAHs), polychlorinated biphenyls (PCBs), and phthalates (PAEs) that are of key interest to urban ecosystems because of their detrimental impacts on human health and long residence times (Wang *et al*. 2013; Quero *et al*. 2015; Kumar *et al*. 2020; Liu *et al*. 2021; Lin *et al*. 2023). Finally, many core urban soil microorganisms that were enriched in ruderal soils are thought to be thermophilic, including Caldisericales, Desulfurellales, and Petrotogales, in addition to having associations with various environmental contaminants. These organisms may therefore be more tolerant to higher temperatures associated with urban heat islands than their mesophilic counterparts (Heaviside, Macintyre and Vardoulakis 2017). The prevalence of these taxa and other taxa with related reported functions in urban soils are suggestive of an urban soil microbiome that may be at least partially adapted to pollutants and extreme temperatures common to urban areas.

When extending our analysis to include urban turf (50% of total abundance contained in turf soils with low corresponding abundance in reference soils), we note an overrepresentation of archaea, along with some of the previously discussed bacterial clades associated with contaminant cycling (e.g., Caulobacterales, Burkholderiales). Archaea may be particularly well-suited to turf soils due to their abilities to thrive in extreme environments. Urban turf is subjected to a unique set of stressors including fertilization, irrigation, mowing, chemical applications for weed control and other uses, and high temperatures from lack of tree canopy cover. Indeed, many of the core archaeal clades enriched in turf are associated with nitrogen cycling (Candidatus Nitrosocaldales, Nitrosopumilales, unclassified Thaumarchaeota), anaerobic metabolisms (Methanopyrales), and/or heat tolerance (unclassified Thermoplasmata, Thermoplasmatales, unclassified Aigarchaeota; Pester, Schleper and Wagner 2011; Hedlund *et al*. 2015; Qin *et al*. 2017; Kallistova *et al*. 2023; Sriaporn *et al*. 2023; Zheng *et al*. 2024).

When considering core functions encoded by the urban soil microbiome, we found a plethora of metal-dependent genes and/or genes related to metal cycling (Table S1). We highlight a suite of genes that were found in all cities, no reference soils, and at least 25% of ruderal soils (Table S2). Many iron-related functions fell in this category including: K05297, rubredoxin---NAD+ reductase, containing a central iron atom, and K07231, a putative iron-regulated protein, K09819, a manganese/iron transport system permease protein. In a study by Goff, et al. (2022), multi-metal contamination led to disruption in iron homeostasis, essential for most microbes, which could be a consequence of interest for urban environments (Andrews, Robinson and Rodríguez-Quiñones 2003). Furthermore, although N cycling genes have been reported previously, we note the dependence of N-cycling genes in the core urban soil microbiome on molybdenum, in particular, *nifK*, *nifD*, and *nifE.* Molybdenum is a known urban contaminant with sources including industrialization and fertilization (Lane *et al*. 2013; Wong *et al*. 2021; Tepanosyan, Yenokyan and Sahakyan 2023). These genes display a functional potential for adaptive response to an anthropogenic environment where greater concentrations of metals and contaminants are present.

When we extended our analysis to genes found in all cities and present in more than 25% of ruderal soils and less than 5% of reference soils, we found further evidence for the role of industrial processes in structuring the urban soil microbiome. Both *phsA* and *psrA*, met this threshold and are involved in thiosulfate reductase, which is important in cycling of metals in the environment (Burns and DiChristina 2009). Specifically, *phsA* is required for hydrogen sulfide production, which is toxic to both humans and the environment (Kourtidis, Kelesis and Petrakakis 2008; Quist and Johnston 2023). Additionally, MerA (K00520 mercuric reductase [EC:1.16.1.1]), encoded by the *mer* operon, was found in 43% of ruderal samples. MerA is a protein encoding a subunit in the mercuric reductase enzyme, allowing organisms to survive in the presence of high mercury (Freedman, Zhu and Barkay 2012). Within the same operon, MerR (K08365), a mercuric resistance operon regulatory protein, was found in ruderal soils at 26% (Boyd and Barkay 2012). Fossil fuel combustion has led to increased mercury pollution in the environment, which is expected to continue to rise, and is also highly toxic to humans and the environment (United Nations 2019; Gworek, Dmuchowski and Baczewska-Dąbrowska 2020; Zhang *et al*. 2021).

We also found a suite of genes associated with fructose and glycerol metabolism, and with cellular infection and replication, in more than 25% of ruderal soils and less than 5% of reference soils. While these genes are central to basic metabolic functions in many microorganisms, their disproportionate presence in urban soils raises questions surrounding additional functions these genes may serve in more polluted environments. Fructose, for example, is commonly implicated in plant-microbiome interactions in agricultural and natural soils (Klepek *et al*. 2010; Sasse, Martinoia and Northen 2018; Chen *et al*. 2023; Tharanath, Upendra and Rajendra 2024). However, the urban environment includes additional sources of fructose, such as food waste enriched in high fructose corn syrup (Duffey and Popkin 2008; Pang *et al*. 2021; Martianto *et al*. 2024; Parsa *et al*. 2024) and industrial byproducts including furfural (Chemical Substances Bureau 2008), that may place a greater importance on fructose cycling in urban soils than more natural soils. We specifically highlight *fruA* (K03332), *frcA* (K10554), *pfp* (K00895), and *hmfD* (K16876), as potential areas for future research. Likewise, glycerol is a major component of manufactured products like fertilizers, soaps, cosmetics, antifreeze solutions, and pharmaceuticals, in addition to its natural occurrences (Gerpen 2005; Hájek and Skopal 2010; Pinto and De Araujo Mota 2014). In particular, *glpT* (K17325), *glpP* (K17322), and *tagE* (K00712), may be disproportionately involved in the cycling of these urban waste products in urban soils versus central metabolic processes. Finally, we elucidate three cellular maintenance genes that could be signatures of heightened viral activity in urban soils – K19123 *casA*, CRISPR system Cascade subunit CasA, K03041, *rpoA1*, DNA-directed RNA polymerase subunit A’ [EC:2.7.7.6], K10726, replicative DNA helicase Mcm [EC:5.6.2.3]. While the exact reason for the enrichment of these genes in urban soils is unknown, soil viral ecology is rapidly progressing to the forefront of microbial ecology due to the potential for soil viral impacts on human health and soil biogeochemistry (Neri *et al*. 2022; Roux and Emerson 2022; Graham *et al*. 2023; Jansson and Wu 2023; Zimmerman *et al*. 2024). The enrichment of putative viral signatures in urban soils further underscores the importance of new investigations into soil viral ecology within urban centers.

Despite many commonalities in urban soil microbiomes, Baltimore soils strikingly contained some unique potential functions that may signify a greater impact of anthropogenic processes on microbial communities in this city. Baltimore’s unique functional potential supported diverse genes associated with metal cycling and aromatic degradation (Table S2). Genes unique to Baltimore include K15725, the *czcCBA* efflux system, which is found to play a role in resistance to cobalt, zinc, and cadmium (Nies 2003), and K04108 4-hydroxybenzoyl-CoA reductase, which is found in anaerobic phenol metabolism for benzene degradation (Breese and Fuchs 1998). We therefore posit that cities that have a relatively recent legacy of industrial pollution and fossil fuel combustion, like Baltimore, may harbor organic contaminants (An *et al*. 2013; Gieg, Fowler and Berdugo-Clavijo 2014; Tan *et al*. 2015) that drive distinct soil functional potential relative to cities with more distant industrial eras or stronger environmental regulations.

## Future Directions

Our research brings to light several exciting avenues for future research into urban ecology and biogeochemistry. Efforts to understand core urban microorganisms and functions will benefit from additional sampling campaigns to capture spatial and temporal fluctuations within microbial communities and from methodologies that describe soil microbiome composition and function in more detail. In particular, transitioning from read-based taxonomy to amplicon-based methodologies, or increasing sequencing depth, will likely improve the breadth and resolution of taxonomic identification, providing deeper insights into the microorganisms driving urban soil biogeochemistry. Investigating microbial gene expression through metatranscriptomics or metaproteomics, particularly focusing on the functions identified in this study, will also deepen our understanding of microbial roles within urban ecosystems. Finally, expanding sample collection to encompass a broader range of cities and intra-city locations, as well as temporal fluctuations, will ensure a more comprehensive and representative dataset. These advances can not only refine current ecological models but also provide a robust foundation for hypothesis-driven research and ecosystem management strategies.

## Conclusion

Studies into the unique organisms and functions contained within the urban soil microbiome are essential as the human population continues to expand. Here, we conducted a re-analysis of 66 urban and reference soil microbiomes using metagenomic sequences from the Global Urban Soil Environment Ecology Network (GLUSEEN). Building upon previous work by Epp Schmidt, et al. (2017, 2019), we revealed a set of core microorganisms associated with the urban soil environment and uncovered a set of potential biogeochemical functions that were overrepresented in urban soils relative to more pristine environments. These included genes associated with industrial byproducts, trace and heavy metal cycling, and glycerol and fructose/fructan metabolism; as well as N-cycling genes previously suggested by Epp Schmidt, et al. (2017, 2019). While we hypothesized that changes in urban soil microbiome taxonomic and functional diversity would be correlated to each other, our results suggested that soil microbial taxonomy was more likely to vary within and across cities than functional potential. Baltimore soils, in particular, had a distinct soil microbiome with functions related to contaminants and heavy metal cycling, that may reflect a more recent history of industrial activity and could be a key insight in maintaining healthy urban ecosystems (Pataki *et al*. 2011). Overall, our synthesis reveals unique and ubiquitous features of the urban soil microbiome that provide a basis for further studies on the effects urbanization has on soil and human health.

## Acknowledgements.

The work was performed by Pacific Northwest National Laboratory, operated by Battelle Memorial Institute for the U.S. Department of Energy under Contract DE-AC05-76RL01830. We thank the Global Urban Soil Environment Ecology Network for providing publicly available data upon which this publication is based. We also thank Caroline G. Simonsen, William L. Peterson, Saige G. Smith, and James Kourum for guidance in drafting this publication.

