## Supplementary figures and images for "Urban soil microbiomes exhibit taxonomic and functional potential for enhanced contaminant cycling"

### Supplemental Figure 1

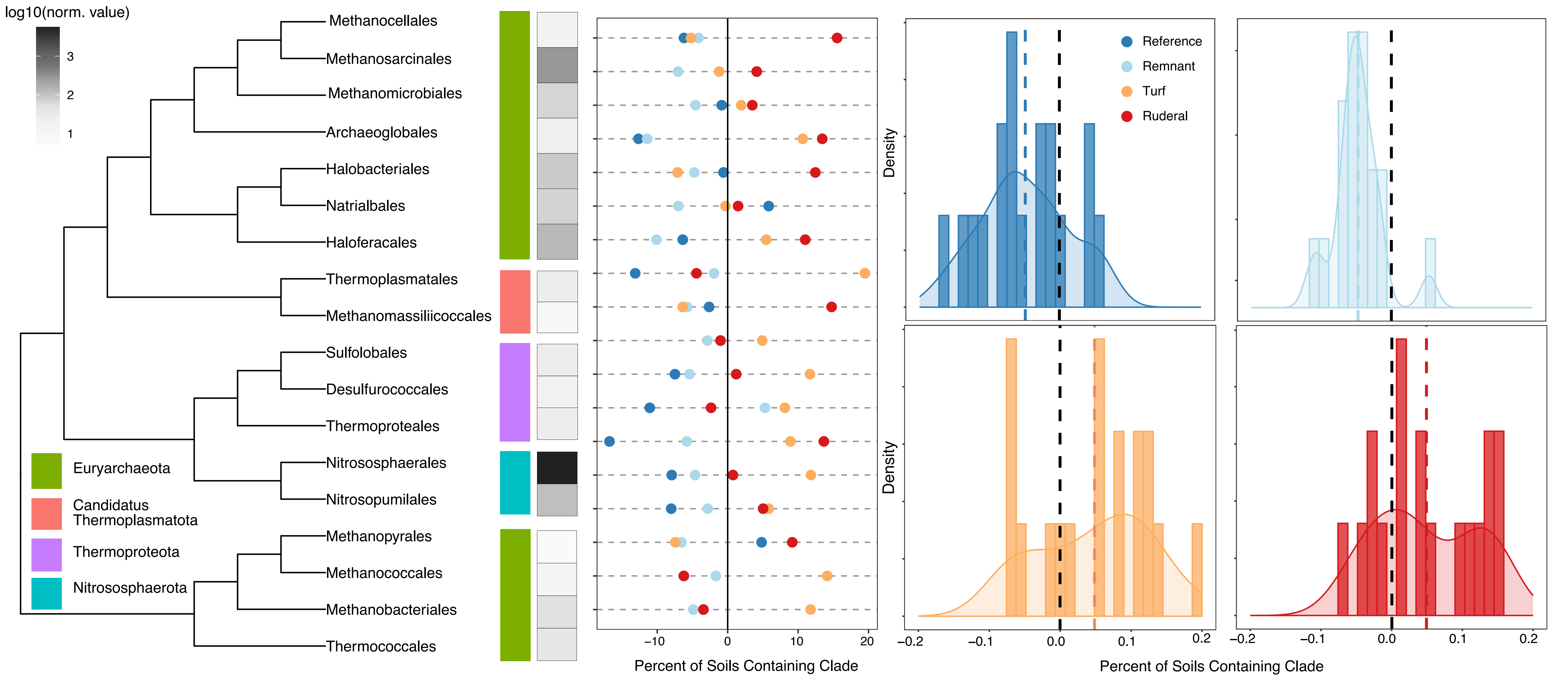
